# Tracking active heterotrophic microbial communities in the Guaymas Basin deep biosphere with BONCAT-FACS

**DOI:** 10.64898/2025.12.16.694778

**Authors:** Andrew Montgomery, Sylvia Nupp, Chase R. Gray, Zackary J. Jay, Virginia Edgcomb, Roland Hatzenpichler

## Abstract

The marine deep biosphere harbors microbial communities that drive organic matter transformations and biogeochemical cycles. Previous work on these communities has focused either on genomic characterization or metabolic activity measurements. However, to understand microbial ecophysiology in the deep biosphere taxonomic identity and metabolic function must be connected on both single-cell and ecosystem scales. In this work, we optimized a bioorthogonal non-canonical amino acid tagging fluorescence-activated cell sorting (BONCAT-FACS) workflow for low-biomass deep-biosphere sediments obtained during International Ocean Discovery Program Expedition 385 (IODP 385). BONCAT-FACS with 16S rRNA gene amplicon sequencing as well as metagenomics of sediment communities was applied to characterize translationally active communities in hydrothermally altered subsurface sediments of the Guaymas Basin. Our results revealed a heterotrophic microbial population throughout all sediments examined, with taxa translationally active down to our deepest sampling point, 154 meters below the seafloor. Based on 16S rRNA gene identities, the translationally active microbial community was dominated by heterotrophic members of the *Gammaproteobacteria*, *Bacilli*, *Deinococci*, and *Alphaproteobacteria*. These taxa are likely key contributors to cycling the large quantities of hydrothermally altered organic matter in Guaymas Basin sediments. To further elucidate the metabolic capacity of active taxa, we mapped 16S rRNA gene amplicons to metagenome assembled genomes (MAGs) previously obtained from IODP 385. These MAGs contained genes associated with C1 metabolism, carbohydrate degradation, and fermentation, indicating that active taxa leverage these metabolisms for energy conservation. Our results demonstrate that BONCAT-FACS provides high-throughput and single-cell insights into the metabolic activity of microbes in the low-biomass marine subsurface.

## Introduction

The marine deep biosphere is one of the largest ecosystems on Earth, supporting an estimated 10^29^ cells [1, 2]. Defined as sediments >1 m below the seafloor, the deep biosphere encompasses a wide spectrum of redox conditions that extend to the energetic and thermal limits of life. Biological and geological transformations of organic matter drive a dynamic, albeit slow, carbon cycle in these sub-seafloor environments [3–6]. While it is clear that microorganisms are metabolically active in the deep biosphere [7, 8], low cell abundances mean the significance of their activity and contributions to biogeochemical cycles remains largely unknown. Direct connections between the taxonomic identity of active microbial populations and metabolic functions are necessary to understand microbial ecology and biogeochemical cycling in the marine deep biosphere.

Uncovering the metabolic capabilities of constituents of microbial communities *in situ* requires a coordinated effort that simultaneously identifies taxa and links them to ecosystem functions. Historically, cultured isolates were required in order to draw conclusions about the metabolic capabilities of specific microbial taxa. However, less than 1% of microbes can be reliably maintained in a laboratory setting [9, 10], and enrichment and isolation conditions can differ substantially from *in situ* conditions, making it difficult to draw conclusions about how microbes live in their natural habitat. Further, cultivation efforts are impeded by low biomass inocula and energy limitations in the deep biosphere that select for microorganisms that grow at extremely slow rates [11, 12]. More recently, culture-independent techniques have been used to elucidate connections between individual microorganisms and their metabolic functions under (near) *in situ* conditions. With the rise of high-throughput sequencing technologies and bioinformatics tools, culture-independent techniques have facilitated studies of the deep biosphere with metagenomics [13–15] to identify microbial taxa, or metatranscriptomics [16–20] to reveal which genes are expressed *in situ*. However, while these approaches offer powerful insights into metabolic potential and community level activity, they are unable to resolve activity at the single cell level and thus cannot link specific taxa with metabolic activity *in situ*.

Bioorthogonal non-canonical amino acid tagging (BONCAT) paired with fluorescence-activated cell sorting (BONCAT-FACS) is a taxonomy-blind Next-Generation Physiology approach [21] that can be used to identify which taxa were translationally active in complex environmental samples during a relatively short incubation without culturing individual organisms [22–26]. BONCAT leverages the incorporation of a synthetic amino acid, such as *L-*homopropargylglycine (HPG), an analog of *L-*methionine, into newly synthesized proteins and its later detection using azide-alkyne click chemistry [22, 27]. Using fluorescence as the mode of detection, FACS can be used to sort translationally active cells and study their identity and genetic make-up via 16 rRNA gene amplicon sequencing [21, 28, 29] or shotgun metagenomics [30] and sorted cells can be subjected to cultivation [22, 31]. BONCAT-FACS has been used to identify active taxa in marine sediments [28], soils [26, 32], the human microbiome [33–35], and other complex environments [29, 36–38]. BONCAT has been compared directly to two gold standards of measuring single cell activity - radioactive isotope probing coupled to microautoradiography [39] and stable isotope probing coupled to Nano-scale secondary ion mass spectrometry [22] - and was found to correlate well with both methods. BONACT coupled to fluorescence microscopy can successfully detect anabolic activity of an *Escherichia coli* culture after incubation with *L*-azidohomoalanine for 2% of its generation time [22] and detect the activity of microbial consortia catalyzing the anaerobic oxidation of methane in deep sea sediments after incubation with HPG for approximately 3.7% of the estimated generation time [9, 10]. Here, we used BONCAT-FACS to identify the translationally active community and, coupled with previously generated community shotgun metagenomics datasets, assess the metabolic capacity of translationally active taxa in the Guaymas Basin deep biosphere.

Guaymas Basin, located in the Gulf of California, is a hydrothermally active rift basin characterized by rapid seafloor spreading and high sedimentation rates. Terrestrial inputs and highly productive waters in the Gulf of California deposit organic matter on the seafloor at rates >1 mm yr^-1^, generating sediment packages that can be hundreds of meters thick [40]. Thermocatalytic alteration of the organic matter, influenced by basaltic sill intrusions and hydrothermal circulation [41, 42], produces a complex mixture of potentially bioavailable organic compounds in the sediment [43]. This includes methane, low-molecular-weight organics, hydrocarbons, and crude oil [44–49]. International Ocean Discovery Program Expedition 385 (IODP 385) drilled the subsurface of the Guaymas Basin to investigate how microbial communities drive organic carbon cycling in the hydrothermally altered deep biosphere. In total, holes were drilled at eight sites (U1545-U1552) to capture unaltered sediment and a spectrum of hydrothermal sediments including extremely hot sediments above a more recently intruded sill-sediment system [50]. To explore the microbial ecology of the Guaymas Basin subsurface, we tracked translationally active taxa in six samples, from six depths at five sites drilled during IODP 385. We were limited in the number of samples available for this work from difficult to access sites, but were able to identify translationally active taxa from these samples. We optimized a previously established BONCAT-FACS workflow with 16S rRNA gene amplicon sequencing to identify translationally active taxa [28, 29] and to assess diversity across biogeochemical gradients. We coupled this with metagenomic analyses from IODP 385 [14] to provide direct insight into the metabolic capacity of active taxa within the Guaymas Basin deep biosphere.

## Materials and Methods

### Sample collection and BONCAT incubation

Sediment cores were collected from Guaymas Basin during IODP expedition 385 onboard drilling vessel *JOIDES Resolution.* Samples were taken at six depths, ranging from 0.76 to 154 mbsf, from five drill holes (Table 1). Intact, round samples were aseptically transferred to the microbiology lab after retrieval on deck and stored at 4°C in heat-sealed gas-tight bags flushed with N_2_ gas to maintain anoxic conditions for up to 30 minutes until processing. Material for metagenome analysis was immediately frozen at −80°C. While we cannot rule out the chance of drilling fluid or laboratory contamination in these samples, extensive efforts to minimize the likelihood of this are described in recent metagenomic analyses performed on the same samples [14, 20]. Briefly, during IDOP 385 a tracer, perfluoromethyldecalin, was introduced during drilling of holes at each site intended for microbiology sampling. Any cores where detectable tracer was measured in the core interior (the portion sampled for BONCAT) were not processed [20]. Working in an anoxic glove chamber, sediments were taken from the interior of the core excluding 2 cm of the core perimeter, homogenized with N_2_-gassed sterile bottom water, and diluted three times to form a slurry. For each sample, four replicate aliquots of 40 or 120 mL, depending on depth (shallow vs. deep), were distributed into Pyrex bottles and sealed with a butyl stopper and aluminum crimp. The volume of aliquots were larger for deeper samples in order to increase the likelihood of detecting enough active cells for sorting. The amino acid analog *L*-homopropargylglycine (HPG; Vector Laboratories, previously Click Chemistry Tools), a synthetic analog of *L-*methionine, was added to three replicates to a final concentration of 50 µM. The fourth replicate, which served as a control, was not amended with HPG. After HPG addition, samples were incubated at *in situ* temperature (Table 1) for seven days.

After seven days, samples were washed with 1x PBS by adding an equal volume of 1x PBS to each sediment slurry. The mixture was briefly vortexed to mix, dispensed to 50ml centrifuge tubes, centrifuged at 3,200 x g for 20 min to pellet the sediment, and the supernatant poured off. A 5mL subsample was frozen in a cryovial at −80°C for whole sediment DNA extraction, and the remaining sediment was amended with 10% glycerol (9:1 sediment to glycerol-TE), and stored at −80°C until further processing. Back at Montana State University, subsequent steps of the BONCAT-FACS workflow including cell extractions, cell sorting, and 16S rRNA gene amplicon sequencing were done following modified protocols [28, 29]. The full details of the workflow are described in the supplementary information.

### Analysis of Metagenome Assembled Genomes

Metagenome assembled genomes (MAGs) were previously generated from IODP 385 [14]. In short, trimmed reads from all drill holes were pooled and co-assembled using MEGAHIT 1.2.9, and MAGs that were at least 50% complete and contained less than 10% contamination were then used for downstream analyses. Taxa appearing in assembled MAGs that also appeared in contamination controls (laboratory extraction blank and a drilling fluid control) were removed from downstream analyses and are listed in Supplementary data of [14]. Taxonomic identities of these MAGs were assigned using GTDB-Tk v2.1.0 [51]. MAG contamination and completeness were checked using CheckM v1.2.2 [52]. Barrnap (v0.9) was used to identify 16S rRNA gene sequences from these MAGs. The 16S rRNA gene sequences of these MAGs, if present, were then compared to the 16S rRNA gene amplicon sequences from sorted samples generated in this study for sequence similarity. From a BLAST nucleotide similarity comparison, we used a cutoff of 90% identify match and 99% for hit length to identify sequences that matched.

The organism identities of these matches were then compared between the MAGs and amplicon sequences to determine if they were identified as the same organism. Importantly, the MAG and 16S rRNA gene amplicon sequences were independently sequenced and taxonomy was assigned using different databases (Silva 138.2 for 16S rRNA gene amplicon sequences, and GTDB-Tk 2.1.0 for MAGs), and taxonomic identification varied between the two. Organism identity was matched based on nomenclature changes in the literature to the best of our ability. For consistency across our datasets, the Silva 138.2 taxonomic identification is used throughout the manuscript. We were able to successfully match six organisms out of the datasets that met our cutoff criteria of 90% for %ID and 99% for %hit, five bacteria and one archaea.

MAGs were annotated with Prokka v1.14.6 [53] separately for archaea and bacteria. Individual genes of interest for the environment were selected based on Dombrowski *et al.* [54]. Carbon metabolism pathways were identified using KEGG BlastKOALA v3.1 [55]. Genes for the Wood-Ljungdahl pathway were identified using an internal protein database generated from cultured methanogens because of the presence of MAGs in the same class as other methanogens.

## Results & Discussion

### Sampling Sites

Samples for BONCAT incubations were collected from drill hole sites, U1545 to U1549, along a transect across Guaymas Basin during IODP Expedition 385 (Table 1). Sites U1545 and U1546 are located ∼52 km off-axis of the central spreading center, and include a relatively unaltered reference (U1545) site and a more thermally altered (U1546) site with deep sediments (∼350-430 mbsf) influenced by a now thermally-equilibrated ancient sill emplacement [50]. Sites U1547 and U1548 were drilled within and adjacent to the much hotter Ringvent site. Ringvent is shaped by a shallow, more recently emplaced sill, exhibits a sill-driven hydrothermal mound, and allows us to compare other samples to a site with higher sediment alteration and elevated temperatures. Site U1549 targeted an off-axis cold seep near a subsurface sill, where methane seepage and hydrates indicate persistent subsurface fluid migration. Sample depths ranged from 0.8 m at site U1546 to 155 m at U1545, and temperatures were between 2.8°C and 46.6°C (Table 1). Sediment and porefluid geochemical data relevant to microbial activity are outlined in Table 1, and comprehensive datasets are available in the IODP 385 expedition reports [56–59].

### Workflow development

We expanded the applications of BONCAT by refining a workflow for detecting translational activity of microbial communities in extremely low-biomass marine deep-biosphere sediments that maximized cell recovery and minimized interference from sediment autofluorescence. To optimize this workflow and test the efficacy of cell extraction protocols, we spiked IODP 385 sediment with *Escherichia coli* (*E. coli*) cells that had been grown with HPG. These sediments had been preserved analogously to the IODP 385 BONCAT samples and incubated without HPG. The presence of BONCAT-positive *E. coli* in cell extracts was used to assess the efficacy of different cell extraction protocols to maximize cell recovery. Additionally, we sought to minimize particulate matter in the final cell suspension. Autofluorescent sediment particles can interfere with FACS analyses when the signal overlaps with the target fluorophore and distorts forward and back scatter, complicating the accurate detection of cells.

The final cell extraction procedure was adapted from an established protocol [28, 60] and consisted of two primary cell separations. The first wash with methanol only was used to extract cells that are more susceptible to lysis from detergents. The second step combined sonication with a detergent mixture (sodium pyrophosphate, Tween80, and methanol) to detach cells from sediment particles. A 50% Nycodenz gradient was used between detergent wash steps to phase-separate detached cells from sediment. Based on the number of cells and amount of autofluorescent sediment in the extracted suspension, we determined that pooling eight technical replicate extracts was sufficient to extract cells from low-biomass samples for downstream processing with FACS. Field samples analyzed in this study had in situ cell abundances that ranged from 10^9^ and 10^8^/cm^3^ (near-surface samples U1546B-1H2 and U1547B-1H2) to 10^4^/cm^3^ (U1547B_8H2 at 65.8 mbsf). Methods for these cell counts and data are shown in [14, 61]. The rapid decline in down-core cell counts in the Guaymas Basin subsurface is consistent with decreasing DNA recovery [61]. We visualized labeling of BONCAT-positive cells from click chemistry in the presort and sorted fractions with fluorescence microscopy (Fig. S1). During method testing and sample analysis, the presort fraction contained cells with and without HPG incorporation, while the sorted fraction only contained HPG-positive cells, demonstrating the ability of our method to successfully sort exclusively BONCAT-positive cells from low-biomass marine deep biosphere sediments.

BONCAT has been successfully used on dozens of bacterial and archaeal phyla in many ecosystems, including the human microbiome [33, 34], marine sediments [23, 62] and water column [63, 64], soil [26], wastewater sludge [65], hot springs [29], the terrestrial subsurface [66] and, more specifically, Guaymas basin sediments [28]. However, to the best of our knowledge, it had yet not been applied to fungi. Initially, we aimed to assess the translational activity of fungi with BONCAT-FACS in this study, so we developed this cell extraction protocol to include fungal cells. We used two fungal isolates from Guaymas Basin that are representative of the fungal community in IODP 385 sediment [67], and tested extraction methods similar to our use of *E.coli* above. Based on our testing, this cell extraction method can be used to separate yeast and filamentous fungal cells from sediments. Furthermore, these fungal strains incorporate HPG in liquid media and can be detected with click chemistry and fluorescence microscopy (Fig. S2), demonstrating that BONCAT workflows can be applied to marine fungi. However, due to issues with downstream processing, we could not reliably detect fungi in the presort and sorted fractions with internal transcribed spacer (ITS) amplicon sequencing. While the exact reason for this issue is unknown, poor extraction efficiency, low fungal biomass (or absence of fungi), the inability to sort certain fungal cells with FACS, or poorly matched PCR primers for the ITS region in natural fungal populations [68, 69] may have all impeded the successful detection of fungi in our samples. Therefore, we chose to exclude any further discussion of fungi from this manuscript and focus on bacteria and archaea inhabiting the Guaymas Basin deep biosphere.

### Diversity of the total, cell extractable, and active prokaryotic community

We combined BONCAT-FACS with 16S rRNA gene amplicon sequencing of sorted cells to identify translationally active bacteria and archaea in the Guaymas Basin deep biosphere. To assess community compositions, the observed diversity (the quantity of unique ASVs) and Shannon’s diversity index were determined for each sample (Fig. 1A-B). For all samples, observed and Shannon’s diversity indices were higher in the whole sediment DNA extract fraction than in the cell-extracted presort and BONCAT-active sorted fractions (Fig. S3). Diversity was generally higher in the shallower sediments, reflecting higher cell counts in those sediments. The two deepest samples, 1547B-8H2 and 1545B-19F3 (66 and 155 mbsf, respectively), exhibited the smallest difference in diversity metrics across all three fractions, especially with respect to the whole sediment DNA extract, taken after incubations but before cell extraction. Compared to shallower samples, a larger portion of the total microbial community was detected in the BONCAT active fraction in these deeper sediments, where there are relatively few cells, carbon is more recalcitrant, energetic conditions are less favorable, and geochemical conditions remain stable over extended timescales [70, 71]. This suggests that the microbial population is highly specialized to fill limited available niches in the deep biosphere.

**Figure 1.**
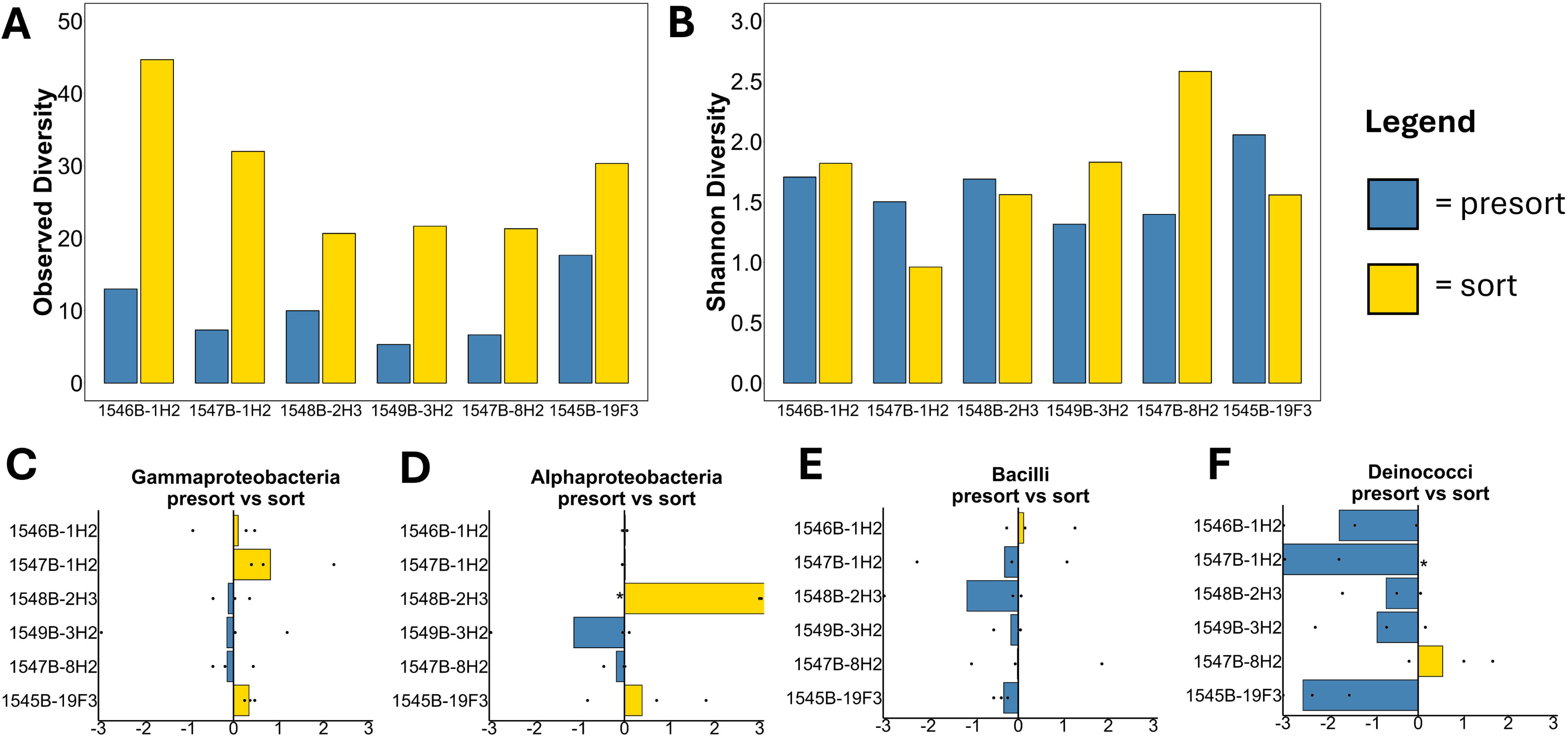
Alpha diversity indices of each sample and fold enrichment changes of the 4 most abundant Classes. Plots of Observed Diversity (A) and Shannon’s diversity index (B) for all sites. Fold enrichment of relative abundance between the pre-sort and sorted fractions for Gammaproteobacteria (C), Alphaproteobacteria (D), Bacilli (E), and Deinococci (F). Fold enrichment was calculated as the ratio of relative abundances of ASVs affiliated with a given class between the pre-sort and sorted fraction, normalized to zero so that equal frequencies equate to zero. In fold-enrichment plots (C-F), yellow bars indicate that a class was more represented in the sorted fraction, and blue bars show that the class was more abundant in the pre-sort fraction. Bars represent the average of triplicate samples, and each point shows the value of each replicate. A * indicated that no ASVs affiliated with that class were detected in either the presort or sorted fraction. Note that samples are presented in order of increasing depth.

To evaluate the changes in diversity metrics between the presort and sorted fractions, we calculated fold enrichment (Fig. 1C-F) between these fractions for the most abundant ASVs, which were affiliated with *Gammaproteobacteria*, *Alphaproteobacteria*, *Bacilli*, and *Deinococci* (Fig. 2 and Table S2). Generally, these abundant taxa were not overly represented in either fraction (enrichment values near zero) or were enriched in the presort fraction (*Deinococci* and *Bacilli*), suggesting that only a portion of those taxa detected in the presort fraction were actually translationally active. In several cases, corresponding ASVs were not detected in either the presort or sorted fraction, which skewed enrichment toward the respective other fraction. For instance, in sample 1548B-2H3, no sequences affiliated with *Alphaproteobacteria* (Fig. 1D) were detected in the presort fraction, artificially inflating the fold enrichment toward the sorted fraction. Similarly, no *Deinococci* affiliated ASVs were observed in the sorted fraction for one replicate from 1547B-1H2 (Fig. 1F). The inability to detect a given ASV in a specific fraction could skew diversity indices, making it difficult to interpret diversity data in those samples.

**Figure 2.**
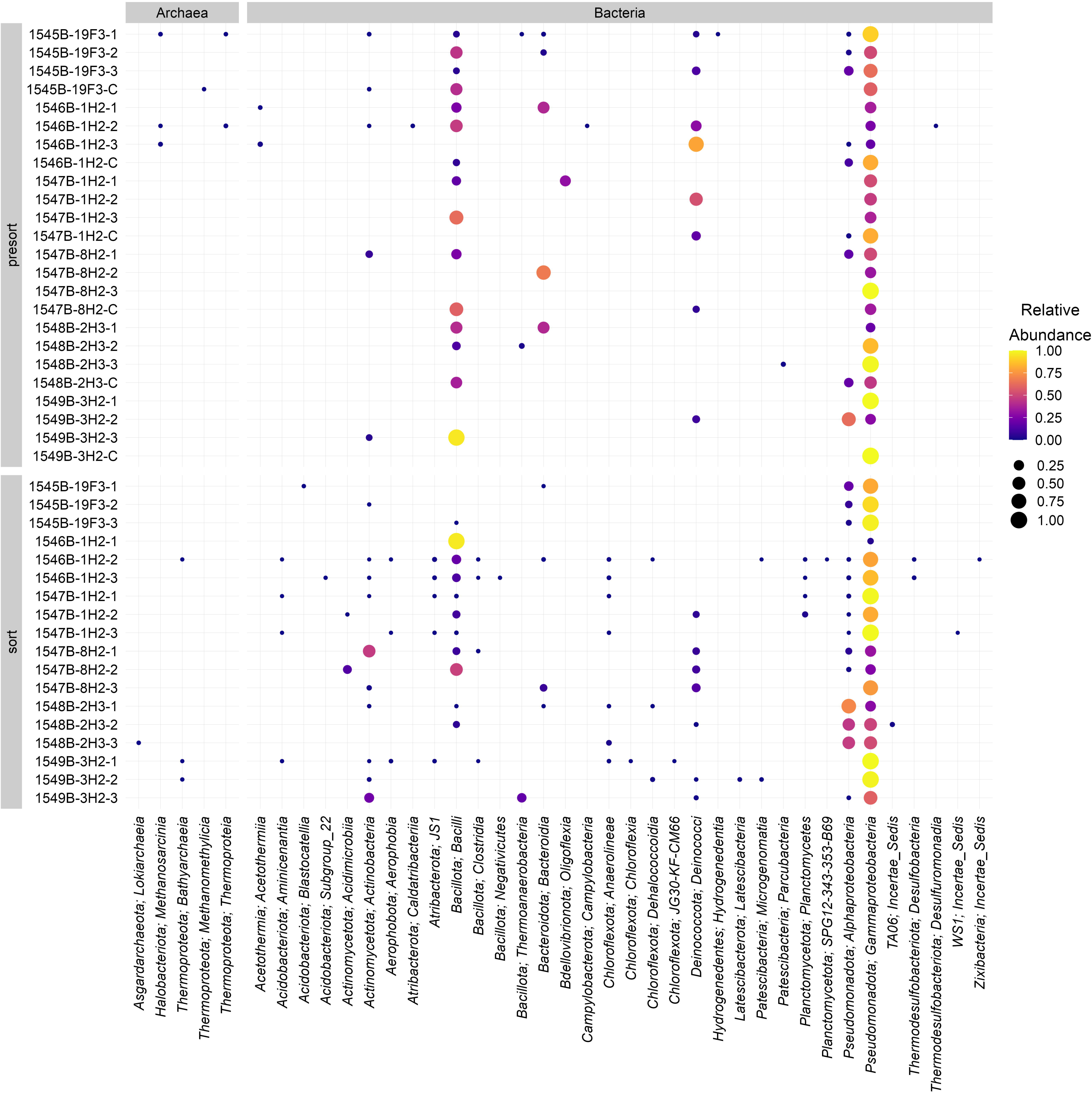
Community composition of cell extract (pre-sort) and BONCAT-positive sorted samples depicted as the relative abundance of ASVs of the 16S rRNA gene collapsed to class level. Fractions are grouped separately on the y-axis. Replicate numbers are indicated by 1-3; C refers to the no-HPG control.

Observed diversity of unique ASVs increased from the presort to the sorted fractions in all samples (Fig. 1A), and Shannon’s diversity increased in 1549B-3H2 and 1547B-8H2 between those fractions (Fig. 1B). It should be expected that diversity would decrease between the presort and the BONCAT active fraction to reflect that only a portion of the community is translationally active and detectable with BONCAT-FACS. One explanation for this outcome is that there were substantially fewer reads in many of the presort samples than in the sorted samples (Fig. S4), likely due to the relatively small subsample (50 µL) taken from the presort samples for DNA extraction and the low amount of biomass in the sample. More ASVs were detected when the number of reads was higher (Fig. S4), which would influence alpha diversity metrics. In the two samples in which Shannon’s diversity increased in the sorted fraction (1547B-8H2 and 1549B-3H2), across all replicates, we detected between three and eleven unique class-level ASVs in the presort samples and between 12 and 30 in the sorted samples. This indicates that sequencing depth directly impacts the detection of additional and/or rare taxa that may have been, in some cases, missed in the presort fraction.

We observed major differences in Shannon’s diversity indices across the BONCAT workflow (DNA extract to presort to sorted fractions). Shannon’s diversity was highest in the DNA extract relative to the presort or sorted fractions for nearly every sample (the only exception was the sorted fraction for 1547B-8H2) (Fig. S3). This is expected for the marine subsurface as extracellular DNA in the sediments may have been amplified with PCR, altering community composition and artificially inflating diversity metrics in the initial DNA extract fraction [26]. Diversity indices were generally lower in the presort and sorted fraction as compared to the DNA extract fraction, as cell extraction for BONCAT-FACS is known to introduce biases [28, 29] that could cause large discrepancies in diversity metrics between the DNA extract and presort fractions. To address this, we optimized the cell extraction procedure for these low-biomass deep biosphere samples, but biased cell recovery is a challenge for any complex sample [21, 72], especially from Guaymas Basin sediments that are enriched with hydrothermally altered organic carbon. Ideally, cell extraction procedures should be tuned to specific samples based on experimental goals and biogeochemical conditions.

### Abundant Taxa and Active Community

To assess the active community within the Guaymas Basin deep biosphere, we analyzed the relative abundances of ASVs in each fraction and compared those frequencies between samples (Fig. 2 and Fig. S5). In the presort cell extract and BONCAT active sorted fractions, we identified a total of 40 bacterial classes and six archaeal classes, which were frequently dominated by a few classes. Across all three fractions (DNA extract, presort, and sorted), ASVs affiliated with *Gammaproteobacteria* were consistently the most abundant taxa detected. *Gammaproteobacteria* had greater than 50% relative abundance within the BONCAT active community in 12 of the sorted fraction samples (Fig. 2). In three of the six samples (1546B-1H2, 1547B-1H2, and 1545B-19F3), all *Gammaproteobacterial* genera were detected in higher relative abundance in the sorted fraction than in the presort fraction (Fig. 2), which indicates members of this group are active in the Guaymas Basin deep biosphere. High abundances of *Gammaproteobacteria* in the BONCAT active fraction likely reflect the heterotrophic nature of this class, which is capable of degrading diverse organic substrates typically found in Guaymas Basin sediments [73–75]. While the relative frequency of *Gammaproteobacteria* was elevated in most samples, differences in relative abundance between fractions could be explained by enrichment of *Gammaproteobacteria* during BONCAT incubation, or methodological biases against other taxonomic groups.

Across all three fractions, we identified 61 genera affiliated with the *Gammaproteobacteria* (Fig. S6 and Table S3) that are associated with diverse heterotrophic metabolisms. As observed for the whole community analysis, the highest number of gammaproteobacterial genera was observed in the DNA extract, and ASVs affiliated with the genus *Thiomicrohabdus* were amongst the most abundant. In contrast, the presort and sorted fractions were primarily dominated by members of *Halomonas,* and to a lesser extent, *Burkholderia-Caaballerionia-Paraburkholderia* and *Acinetobacter*, which had the highest relative frequency in the deepest sample (1545B-19F3). In several samples, ASVs belonging to *Halomonas* (1546B-1H2, 1547B-1H2, and 1549B-3H2) and *Acinetobacter* (1545B-19F3) exhibited a relative frequency >75% in the BONCAT active fraction in at least two replicates. This was a sharp increase compared to the other fractions which, across all of those samples, were typically less than 20% for the DNA extract and all less than 3% for the presort fractions. The relative abundance of *Halomonas* demonstrated the largest observed increase between the presort and sorted fractions (Fig. S6). In particular, BONCAT active *Halomonas* ASVs were detected at elevated frequencies in samples with temperatures <15°C (Table 1), compared to warmer samples (∼38°C). Sediment temperatures can fluctuate by >50°C on relatively small temporal and spatial scales due to hydrothermal activity [76], which may create a selective pressure on the microbial community. This indicates that some members of the *Halomonas* genus remain metabolically active at lower temperatures and withstand large temperature changes characteristic of hydrothermal sediments. Genomic analyses of these communities revealed genes associated with central carbon metabolism, CAZymes, hydrocarbon degradation, and nitrogen, sulfur, and arsenic cycling, suggesting that *Gammaproteobacteria* are key drivers of organic matter remineralization and redox transformations in the subsurface [75, 77].

In addition to *Gammaproteobacteria*, abundant ASVs in the sorted fraction were affiliated with *Alphaproteobacteria* throughout our samples. The relative frequency of translationally active *Alphaproteobacteria* was between 28-52% for three replicates in the relatively warm (∼14°C) sediment sample from 1548B-2H3 and up to 19% in the deepest sample from 155 mbsf at the more thermally moderate site 1545B (∼39°C). After BONCAT incubation, *Alphaproteobacteria* exhibited higher relative frequencies in the sorted fraction in 4 samples compared to the presort fraction (Fig. 1D). Some of the supposed increase in frequency between fractions was due to low detection of *Alphaproteobacteria* ASVs in the presort fraction. Regardless, the presence of *Alphaproteobacteria* in the sorted fractions demonstrates that members of this class are active in the Guaymas Basin deep biosphere. Similar to our observations for the *Gammaproteobacteria*, this is consistent with previous studies that have demonstrated high abundances of *Alphaproteobacteria* and associated heterotrophic activity in the marine deep biosphere [73, 78–80].

Members of the class *Deinococci* were detected in all fractions, and consistently exhibited the highest relative abundance in the presort fraction compared to the DNA extract and active sorted fractions (Fig. 2 and Fig. S5). At Ringvent, in sample 1547B-1H2 (2.2 mbsf), *Deinococci* ASVs constituted up to 54% of the presort fraction, compared to less than 1% in the DNA extract. This higher frequency in the presort fraction suggests our cell extraction method is biased in favor of *Deinococci* cells, which may influence relative abundances in the BONCAT active fraction. In sample 1547B-8H2 (66 mbsf), members of this class accounted for up to 15% relative abundance in the BONCAT active fraction and were less than 6% in the corresponding presort fraction. In hydrothermally impacted habitats, *Deinococci*-affiliated sequences have been detected via 16S rRNA gene amplicon survey [81]. Additionally, analysis of 52 MAGs assigned to *Deinococci* from IODP 360 to Atlantis Bank revealed genes associated with broad anaerobic and heterotrophic metabolisms, including fatty acid and aromatic compound degradation (*e.g.*, acyl-CoA dehydrogenase and benzoyl-CoA), acetogenesis, nitrogen and sulfur cycling, and metal reduction [82]. This versatility can promote *Deinococci* metabolism across the dynamic geochemical gradients observed in the Guaymas Basin deep biosphere (Table 1).

A total of 7 classes affiliated with phylum *Firmicutes* were detected throughout our samples, and the most abundant ASVs from this phylum were assigned to the *Bacilli* class in the presort and active sorted fractions. Although ASVs affiliated with *Bacilli* were detected throughout the sediment, the highest relative abundances in the BONCAT sorted fraction, up to ∼20% (one replicate was above 95%), were observed in the shallower samples (1546B-1H2 and 1547B-1H2) (Fig. 2). In those shallower depths, above the sulfate methane transition zone, where sulfate is the dominant terminal electron acceptor (Table 1) for anaerobic metabolisms, BONCAT-active ASVs affiliated with *Firmicutes* may couple sulfate reduction to organic matter degradation [83]. *Firmicutes* isolates cultured from several marine deep biosphere sites degrade a broad suite of organic substrates, including acetate, polysaccharides, amino acids, alcohols, nucleotides, and microbial necromass, indicating a key role in heterotrophic carbon cycling [28, 73, 84, 85]. This metabolic capacity was substantiated by metagenomic analysis that also demonstrated high thermal tolerance, exceeding 80°C [86]. Additionally, members of the *Firmicutes* phylum within the *Clostridia* class were associated with degradation of crude oil and polycyclic aromatic hydrocarbon degradation in methanogenic enrichments [87]. Hydrothermally induced pyrolysis of organic matter in Guaymas Basin sediments generates petroleum hydrocarbons that are dispersed throughout the sediment column and can fuel diverse microbial metabolisms [45, 46, 88]. In the Guaymas Basin specifically, *Firmicutes* have been linked to both sulfate and nitrate reduction [83] and have dominated enrichment cultures amended with methanol, monomethylamine, and ammonium [89]. Given the wide metabolic capabilities of the *Firmicutes* and the high frequency in the BONCAT active fraction, members of this phylum likely contribute significantly to organic matter degradation and elemental cycling in Guaymas Basin sediments.

### Implications for deep biosphere carbon cycling

The BONCAT active fractions were dominated by a heterotrophic microbial population capable of metabolizing a wide range of organic carbon in the Guaymas Basin deep biosphere. To further assess this metabolic capacity and explore potential implications for biogeochemical cycling in hydrothermal sediments, we mapped 16S rRNA gene ASVs from the BONCAT active fraction to annotated metagenome assembled genomes (MAGs) from IODP 385 [14]. Six unique MAGs associated with the BONCAT active fraction were identified as members of 5 bacterial phyla (*Planctomycetota*, *Chloroflexota*, *Acidobacteriota*, *Aerophobota*, and *TA06*) and 1 archaeal phylum (*Thermoproteota*). These MAGs were 52.7-95.9% complete and all had contamination below 10% (Table S4).

The translationally active MAGs harbored genes associated with diverse carbon cycling pathways, including carbohydrate degradation, C1 metabolism, and fermentation, plus nitrogen and sulfur cycling (Fig. 3). MAG_005 (95.91% complete, 9.60% contamination) was assigned to the phylum *Planctomycetota* and encodes complete pathways for glycolysis and galactose degradation, and a nitrite reductase gene. Recently, the first anaerobic culture of *Planctomycetota* isolated from marine sediments experimentally demonstrated functional capacity for these metabolisms [90], validating the genomic potential of that isolate and MAG_005. MAG_054 (52.66% complete, 3.54% contamination), assigned to the class *Anaerolineae*, contained genes associated with carbohydrate degradation and fermentation, both of which have been demonstrated by cultured *Anaerolineae* [91]. Similarly, NanoSIMS experiments implicated diverse *Chloroflexota* (phylum of *Anaerolineae*) in fermentation and broad heterotrophic metabolisms in hypersaline photoheterotrophic mats [92]. In their experiments, Lee *et al*. (2014) ascertained ^13^C-substrate assimilation by *Chloroflexota* members using phylum-specific FISH probes, but could not resolve taxonomic affiliations on class level or below, or implicate other phyla in carbon cycling [92]. In contrast, BONCAT-FACS workflows provide a high-throughput community-level analysis of active taxa, and when coupled with metagenomics, support the investigation of the metabolic potential of sorted cells [66]. Taken together, BONCAT-FACS, paired with metagenomics, can corroborate the results from culture-dependent and -independent approaches, while also providing metabolic insights at the community level.

**Figure 3.**
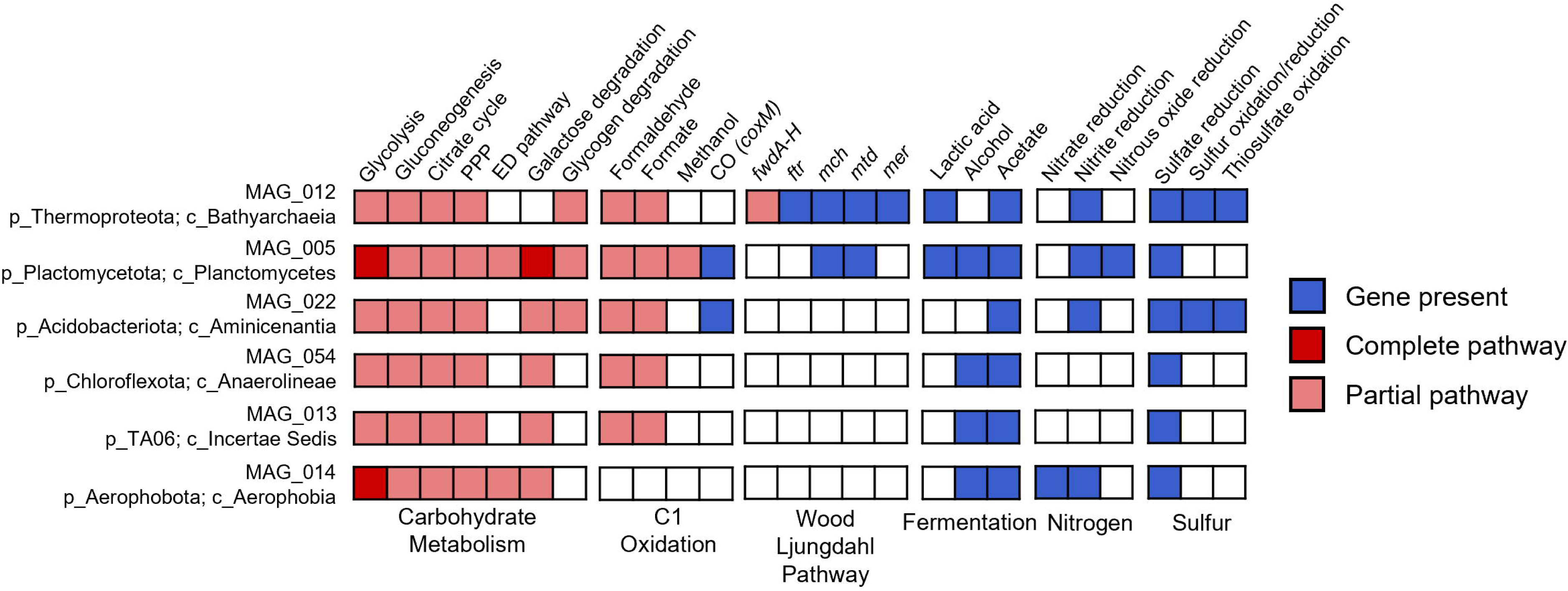
Heat map of metabolic potential of the six translationally active MAGs. A subset of genes was assessed. Squares indicate the presence of a functional gene (blue), complete metabolic pathway (red), or partial metabolic pathway (pink). White squares indicate that no gene(s) affiliated with that pathway were detected. MAGs are numbered following identities in (Mara et al., 2023). Abbreviations: PPP (pentose phosphate pathway), ED pathway (Entner-Doudoroff pathway). MAG_012 (84.65% complete, 9.60% contamination), MAG_005 (95.91% complete, 7.52% contamination), MAG_022 (87.01% complete, 4.23% contamination), MAG_054 (52.66% complete, 3.54% contamination), MAG_013 (86.40% complete, 0.78% contamination), MAG_014 (88.80% complete, 8.25% contamination).

Most importantly, metagenomic analysis, corroborated by 16S rRNA gene identities, revealed a diverse, translationally active microbial community in the Guaymas Basin deep biosphere with the metabolic capacity for a range of carbon transformations and broad anaerobic metabolisms. Our results confirm that these taxa are active, and that under the right biogeochemical conditions (*e.g.,* substrate availability, temperature, etc.), the organisms represented by these MAGs likely utilize these metabolisms for energy conservation in the deep biosphere. However, experimental evidence is necessary to confirm which of these metabolisms are used *in situ* under different conditions. Additionally, the specific ASVs affiliated with these MAGs were not necessarily amongst the most abundant or widespread taxa in the BONCAT sorted fraction. Thus, the extent to which these microbes drive carbon transformations is yet unknown. Future experiments that combine BONCAT-FACS with substrate amendment [28, 29] and paired metagenomics [66] and/or metatranscriptomics could provide deeper, community-level insights into the metabolic capabilities of translationally active taxa in the marine deep biosphere.

## Conclusion

In this study, we assessed the total, cell-extracted (presort), and translationally active (sorted) microbial community in contrasting anoxic sediments of the Guaymas Basin deep biosphere. We identified a diverse extant microbial community down to 154 meters below the sediment surface, and the DNA extract contained 139 unique classes. We optimized our cell extraction and FACS procedures for low-biomass deep biosphere samples and successfully identified 46 classes in the presort and sorted fractions. Alpha diversity metrics decreased sharply from the DNA extract fraction to the presort fraction, likely due to challenges associated with separating certain taxa from the sediments, but were comparable between the presort and sorted fractions.. *Gammaproteobacteria* overwhelmingly dominated the BONCAT-active fraction, followed by *Alphaproteobacteria*, *Firmicutes*, and *Deinococci*. These phyla have the capacity for a wide range of heterotrophic metabolisms and drive microbial-mediated recycling of hydrothermally altered organic matter in the Guaymas Basin deep biosphere.

We demonstrated that BONCAT-FACS can be used to assess microbial activity in low-biomass marine sediments on a community level. Additionally, we paired this with metagenomic analysis from IODP 385 to identify six translationally active MAGs and assess their metabolic capacity. BONCAT-FACS coupled metagenomics links single-cell activity and genetic potential to understand community-level dynamics in a high-throughput manner [21, 66]. This approach could be further optimized in the future to generate metagenomes directly from the sorted fraction and/or by combining experimental amendments with whole sediment metagenomics to more tightly link taxa with metabolic function. Importantly, our workflow, while optimized for low-biomass samples, was likely biased against rare taxa, given the decrease in diversity between fractions. These biases could be exacerbated if combined with metagenomics, which would require greater quantities of biomass for FACS and sequencing compared to gene amplicon sequencing. We were limited in this study by the difficulty of accessing the study site, resulting in a small number of samples. However, we were still able to apply this methodology to deep biosphere samples successfully. With careful benchmarking and optimization, BONCAT-FACS and metagenomics, when combined, offer a powerful approach for connecting microbial taxa and metabolic function in the marine deep biosphere and other complex environmental samples.

## Acknowledgments

We would like to thank the scientific party and D/V JOIDES Resolution crew from IODP 385 for their assistance in sample collection. Amplicon sequencing was performed by the Molecular Research Core Facility at Idaho State University (RRID:SCR_012598). This work was supported by NSF funding to A.M. (PRFB 2010880), R.H. (OCE-2049445), R.H. and V.E. (OCE-2046056), V.E. and R.H. (OCE-2046799), and V.E. (OCE-1829903). Large language models (LLMs) were used to assist in developing data visualization scripts.

## Author Contributions

R.H. and V.E. conceptualized and supervised this study. V.E. collected the samples and performed shipboard sample processing for BONCAT incubations. A.M., S.N., and C.G. developed the methodology, performed experiments, and curated data. A.M., S.N., and Z.J. performed bioinformatics analysis. A.M. and S.N. analyzed the data and wrote the manuscript. All authors edited and gave critical feedback on the final manuscript. A.M., V.E., and R.H. raised funding. A.M. and S.N. contributed to this work equally.

## Data Availability

Amplicon sequence data generated for this manuscript are publicly available under NCBI Sequence Read Archive (SRA) Bio Project accession number PRJNA1356905 (https://www.ncbi.nlm.nih.gov/bioproject/?term=PRJNA1356905) and reads are available under accession numbers SRR35956508-SRR35956585. The metagenome assembled genomes are publicly available under NCBI SRA Bio Project accession number PRJNA909197 (https://www.ncbi.nlm.nih.gov/bioproject/?term=PRJNA909197), and reads are under accession numbers SRR22580794-SRR22580807 and SRR23614663-SRR23614677. The full biogeochemical datasets from IODP 385, including data presented in this manuscript, are available in the IODP expedition report (http://publications.iodp.org/proceedings/385/385title.html).

