## Supplementary figures and images for "Tracking active heterotrophic microbial communities in the Guaymas Basin deep biosphere with BONCAT-FACS"

### Figure S1

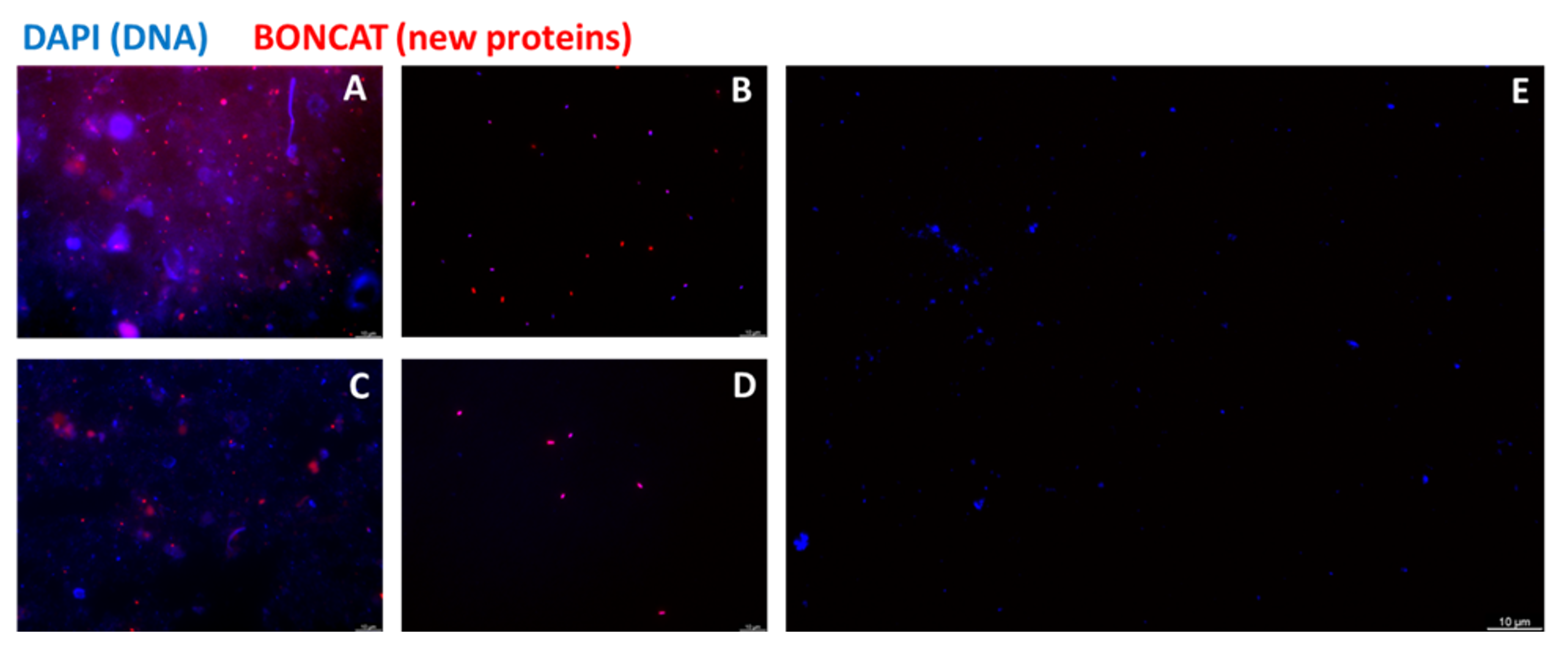

### Figure S2

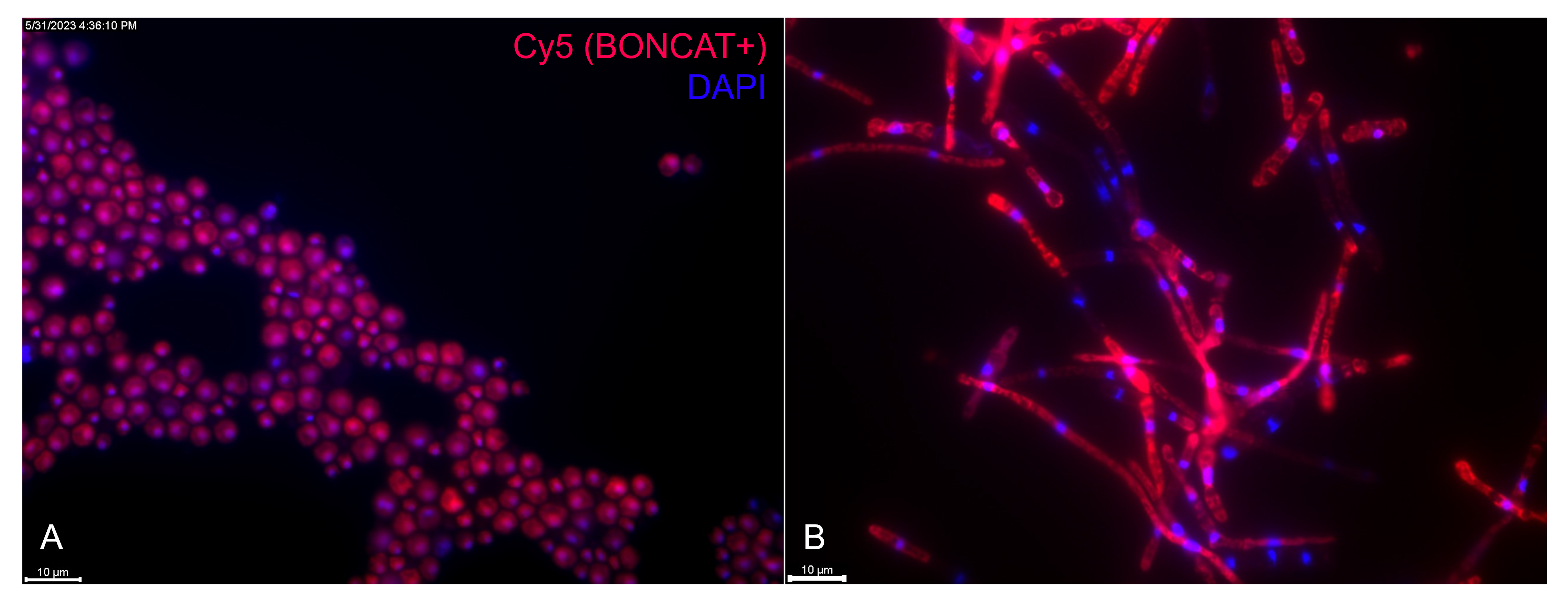

### Figure S3

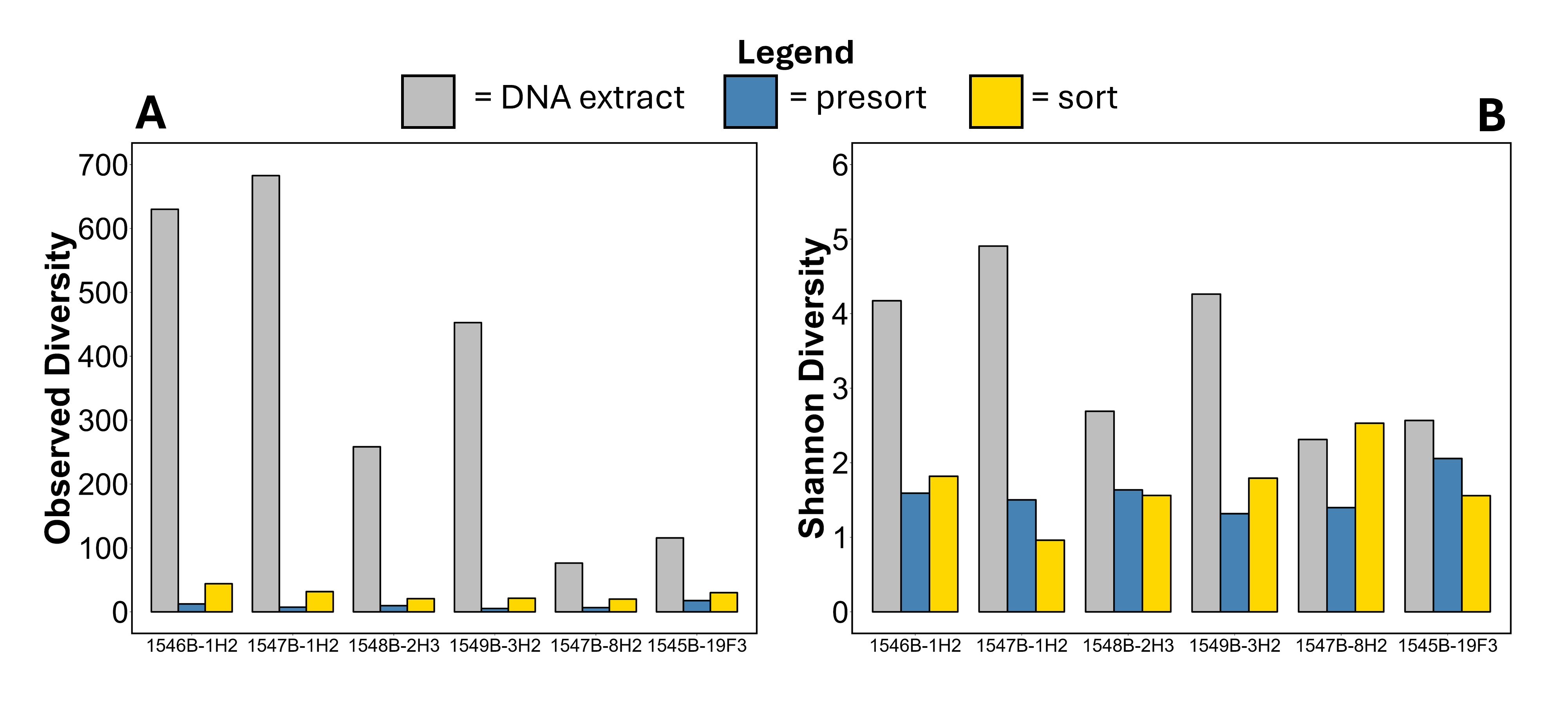

### Figure S4

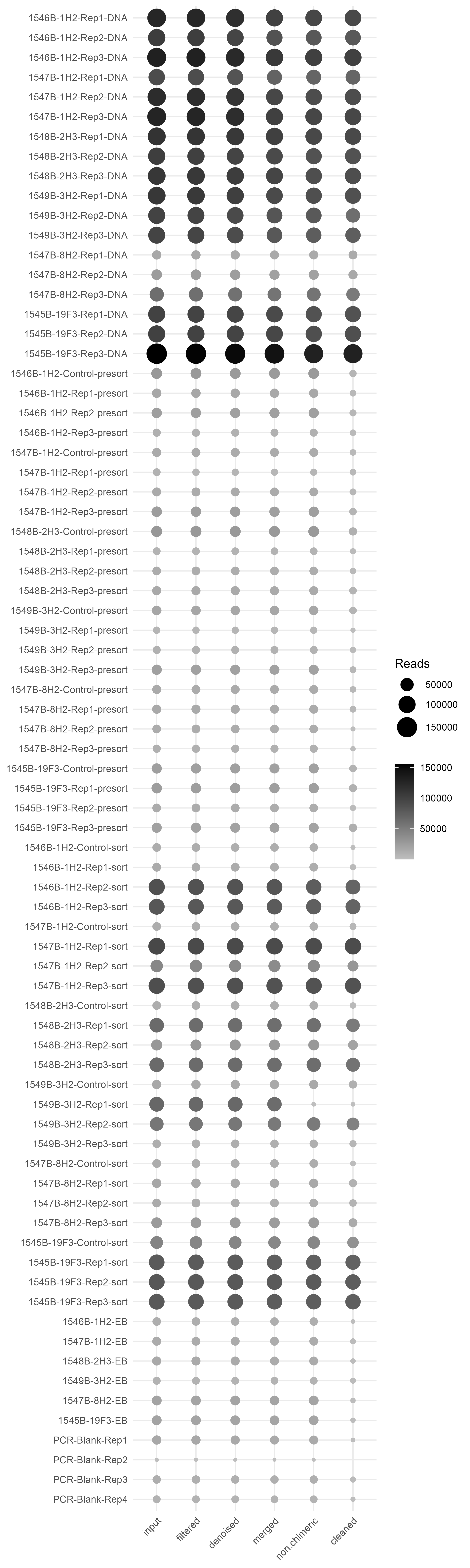

### Figure S5

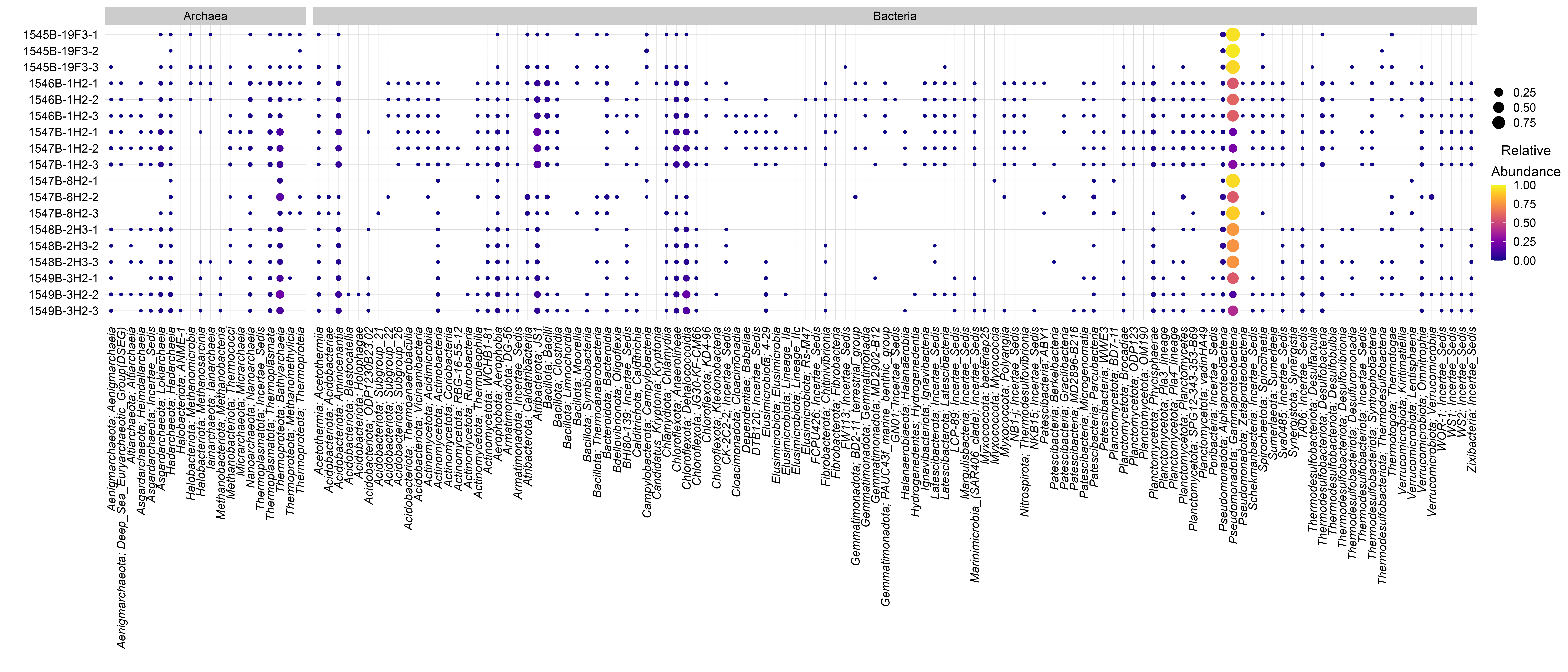

### Figure S6

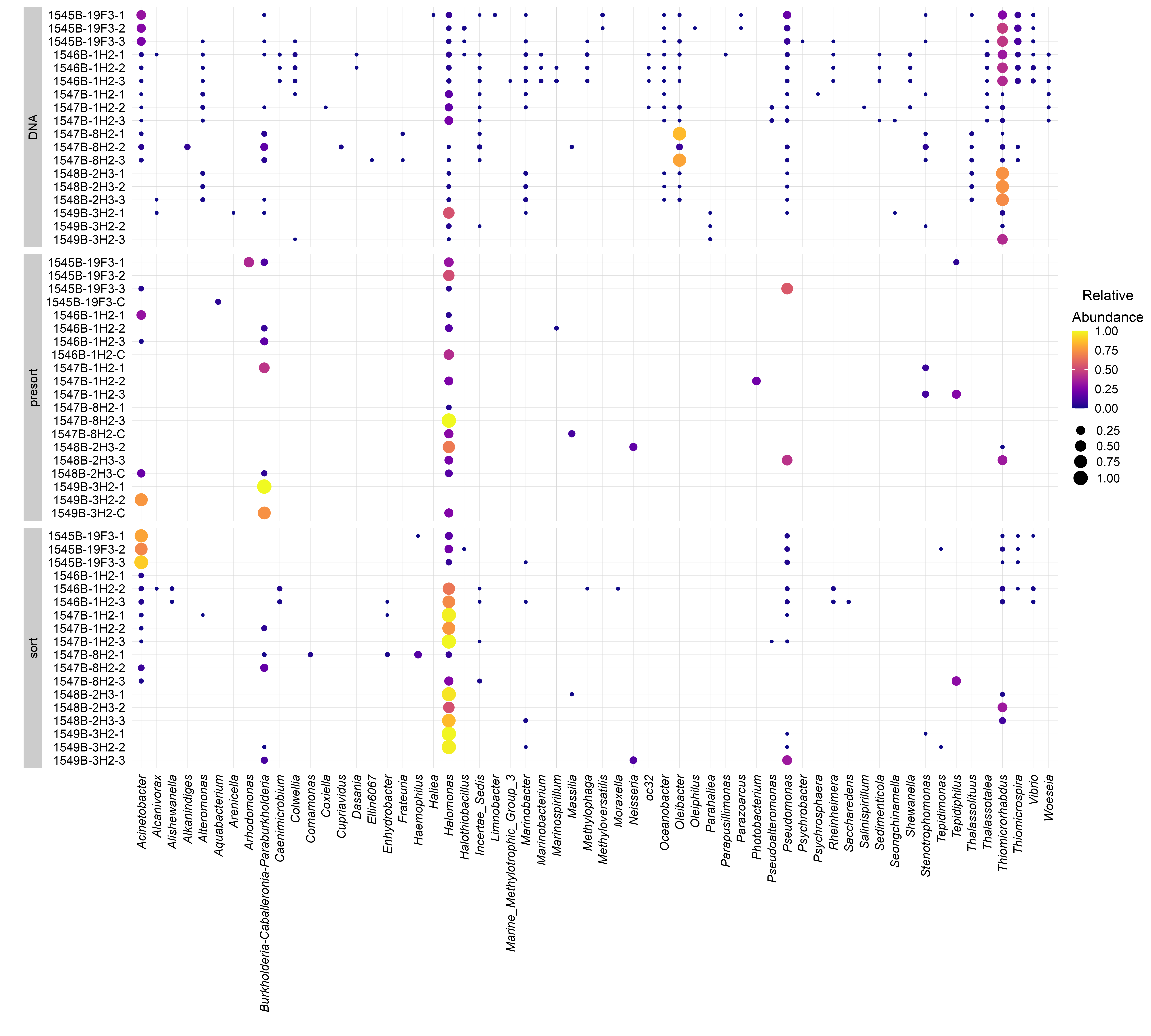

### Figure S7

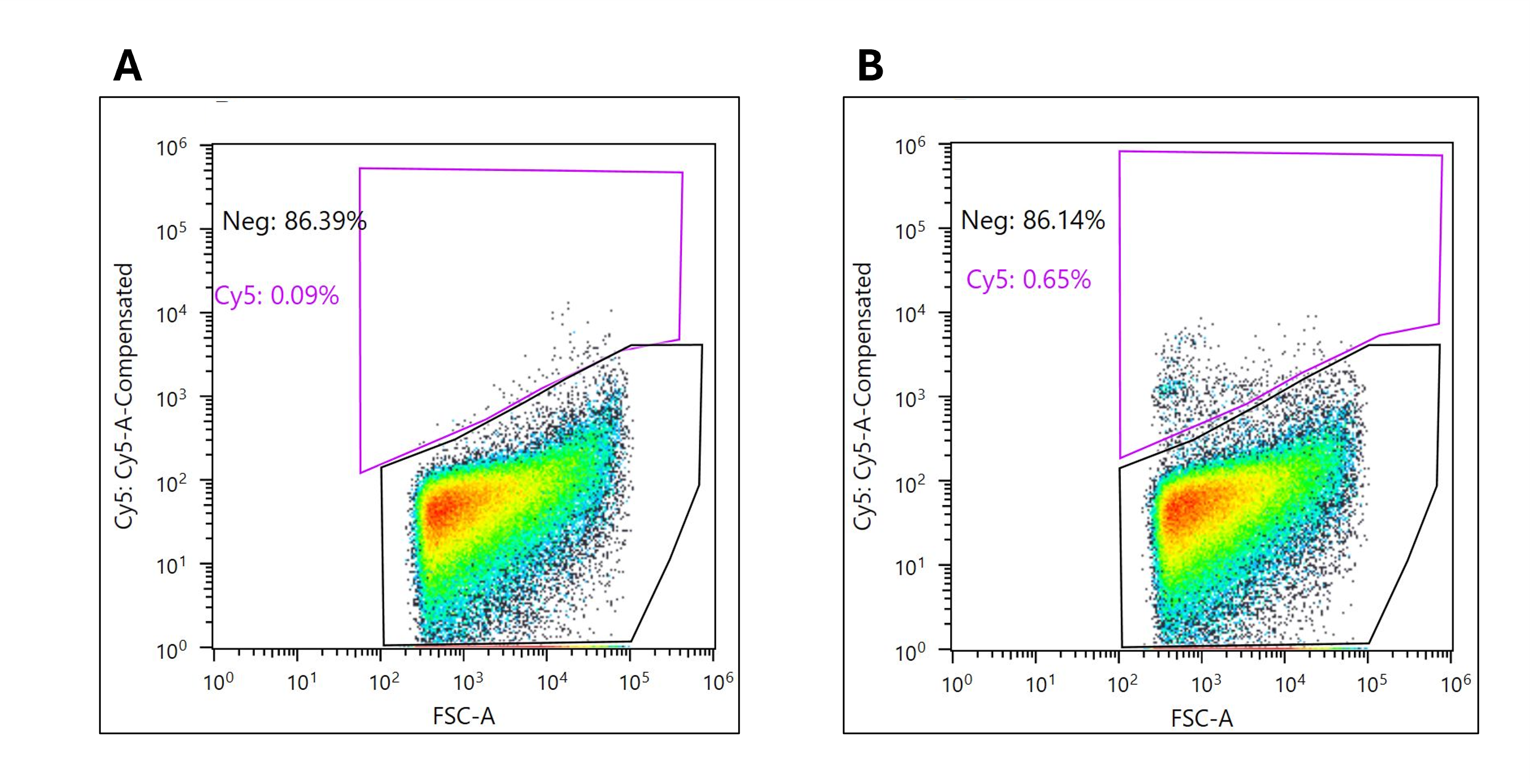
