## Supplementary material for "Tracking active heterotrophic microbial communities in the Guaymas Basin deep biosphere with BONCAT-FACS": SI Figures and Tables

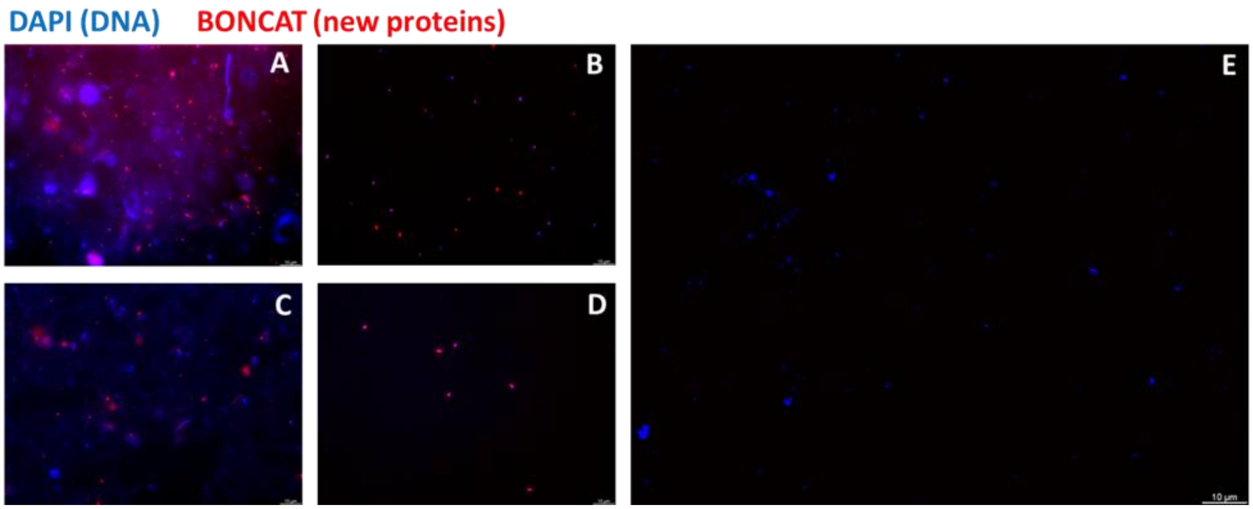


**Figure S1**. Fluorescence microscopy images of IODP 385 BONCAT samples. Images from 1545B-19F3 Replicate A pre-sort (A) and sorted (B), 1548B-3H2 Replicate B pre-sort (C) and sorted (D), and 1548B-3H2 no HPG control pre-sort (E). All cells are stained with the general nucleic acid stain 4′,6-diamidino-2-phenylindole (DAPI) in blue. Cells that incorporated HPG were stained with Cy5 using click chemistry and are depicted as red. Note that the pre-sort samples (A and C) contain cells with and without HPG incorporation, while sorted samples only contain cells that had incorporated HPG (red). Some sediment leftover from cell extraction appears blue, either from autofluorescence or non-specific DAPI staining. Scale bars depicted on each image are 10 μm.

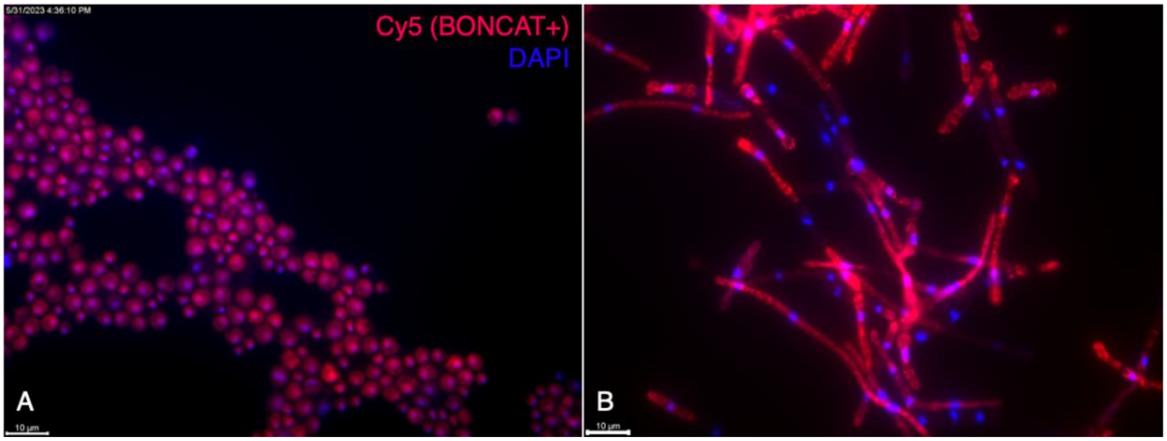


**Figure S2**. Fluorescence microscopy image of fungal isolates showing successful BONCAT labeling. *Rhodotorula sp.* (A) and *Cadophora sp.* (B) were grown with HPG in liquid media, preserved with GlyTE, and subjected to click chemistry to identify HPG incorporation. The red stain indicates BONCAT-positive cells labelled with Cy5, and the blue stain represents the general DNA stain DAPI. The scale bar is 10 μm.

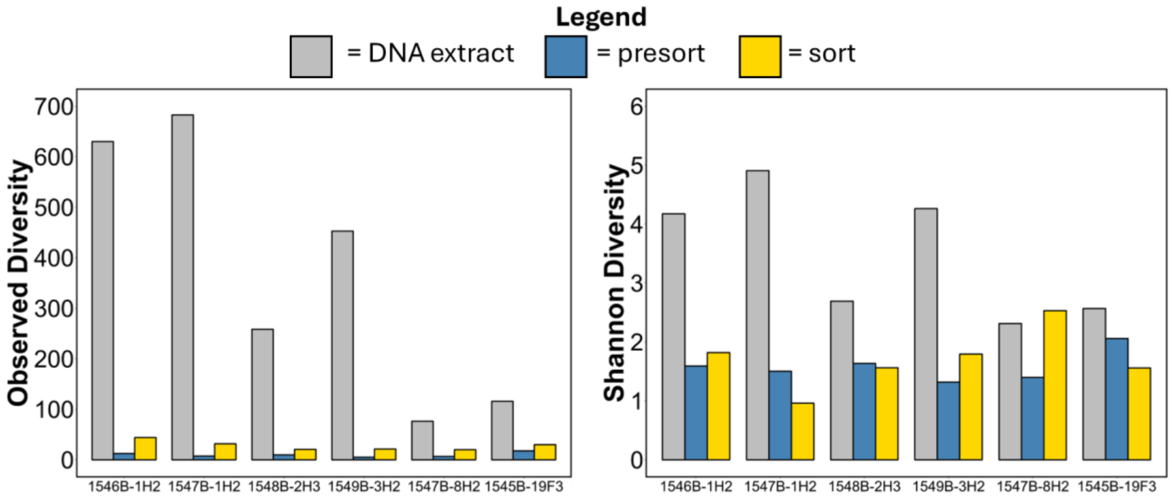


**Figure S3.** Alpha diversity indices of each sample. Observed diversity (A) and Shannon's diversity (B) for DNA extract, cell extracted pre-sort, and BONCAT-sorted fractions from all samples.

**Figure S4.** Sequence processing for 16S rRNA gene amplicons. Samples are grouped by fraction (DNA extract, cell extracted fraction/presort, sorted, and controls) and depth. The size and color of points reflect the number of reads at each processing step. Cleaned = number of reads after processing and removal of contaminant sequences. Control replicates for presort and sort samples are the no-HPG controls. Extraction blanks (EB) were extracted alongside samples and then treated as presort samples. Sorted control replicates, extraction blanks, and PCR blanks were used to assess contaminants and were removed from the dataset prior to community analysis.

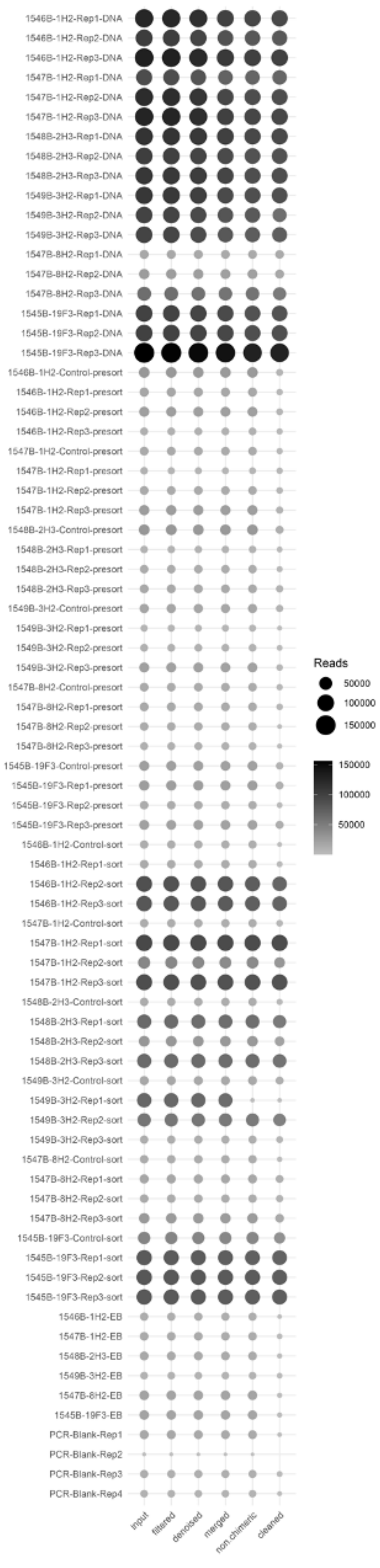

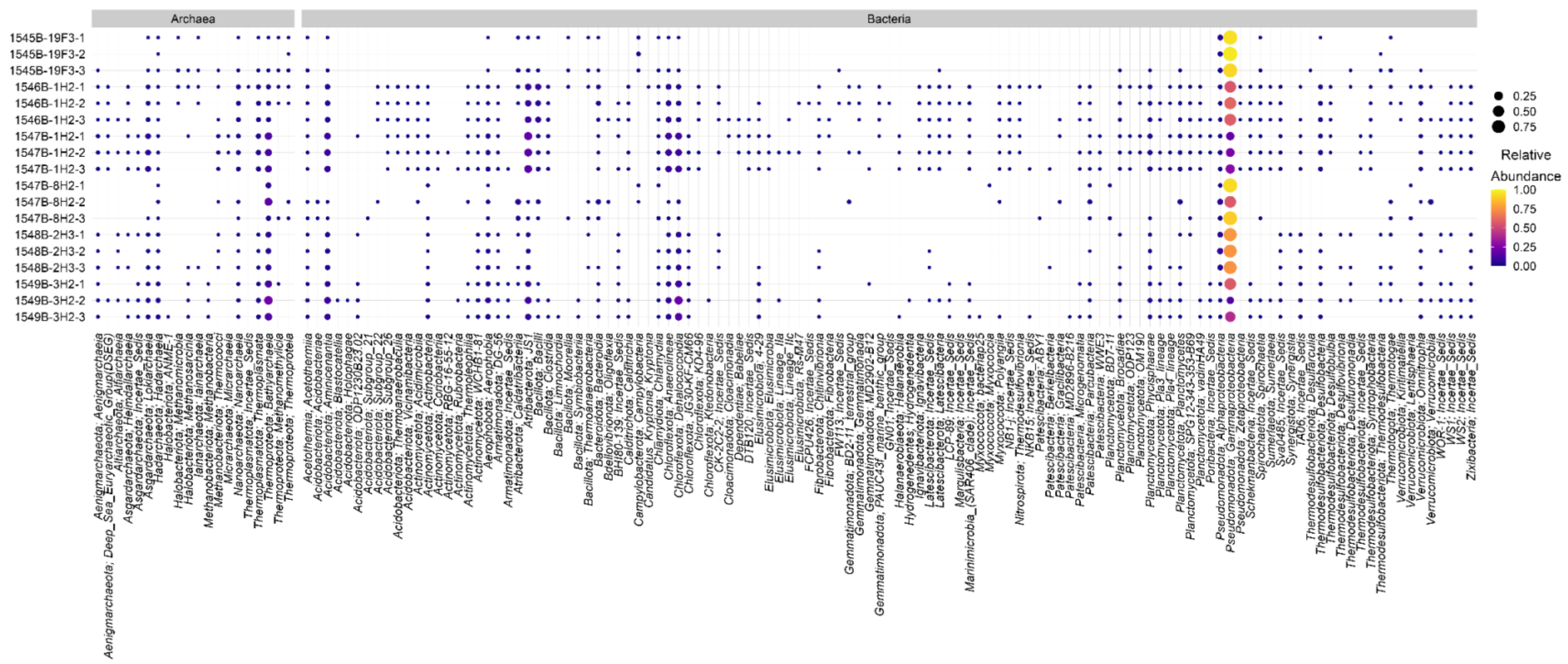


**Figure S5**. Relative abundance of ASVs from 16S rRNA gene amplicon sequencing from whole sediment DNA extracts. Replicates are denoted Rep1-3.

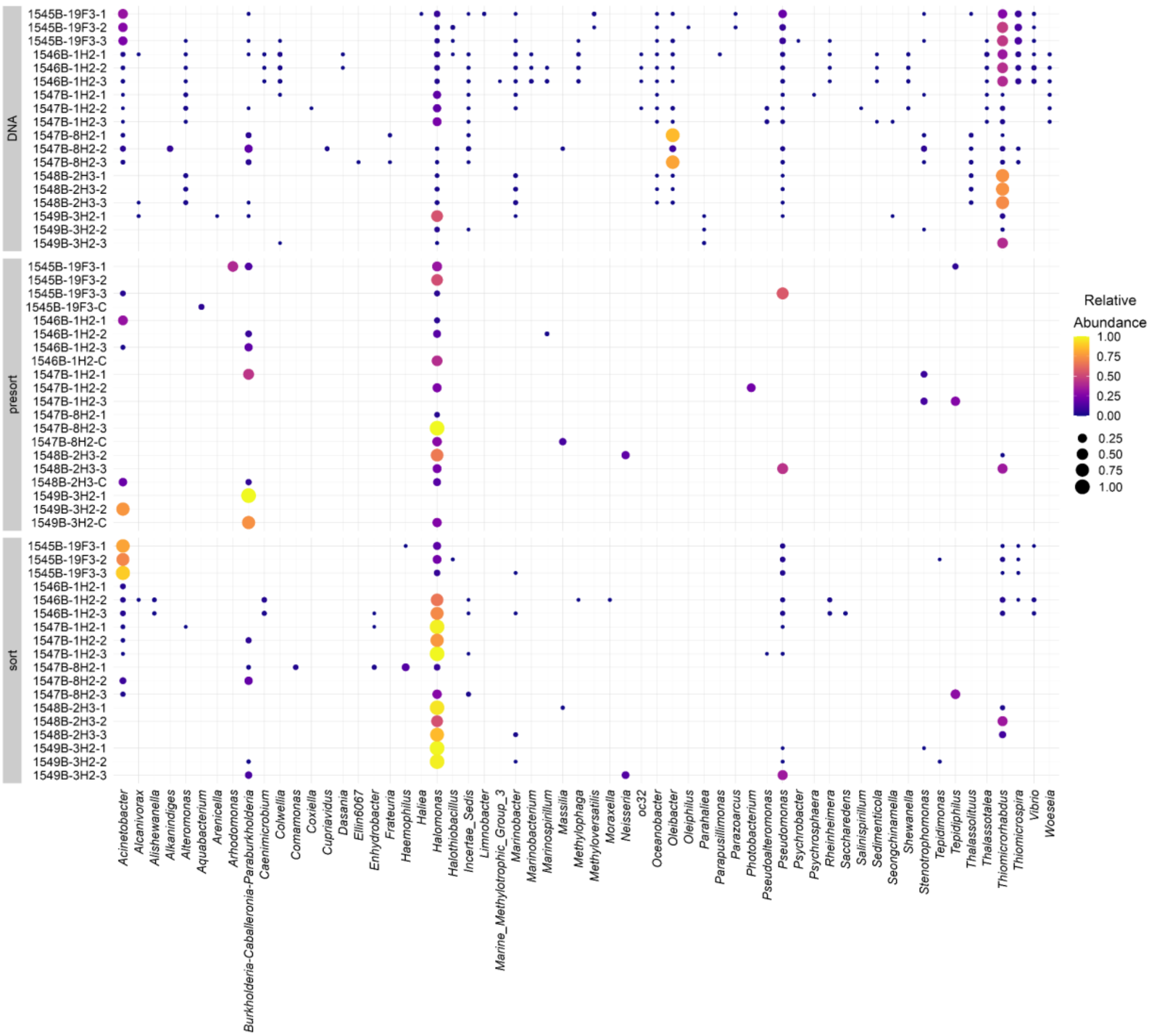


**Figure S6**. Relative abundance of genus-level ASVs, affiliated with the Gammaproteobacteria, from 16S rRNA gene amplicon sequencing for each fraction. Fractions are faceted on the y-axis for the DNA extract (DNA), cell extract (presort), and BONCAT positive fractions (sort). Replicate numbers are 1-3; C refers to the no-HPG control.

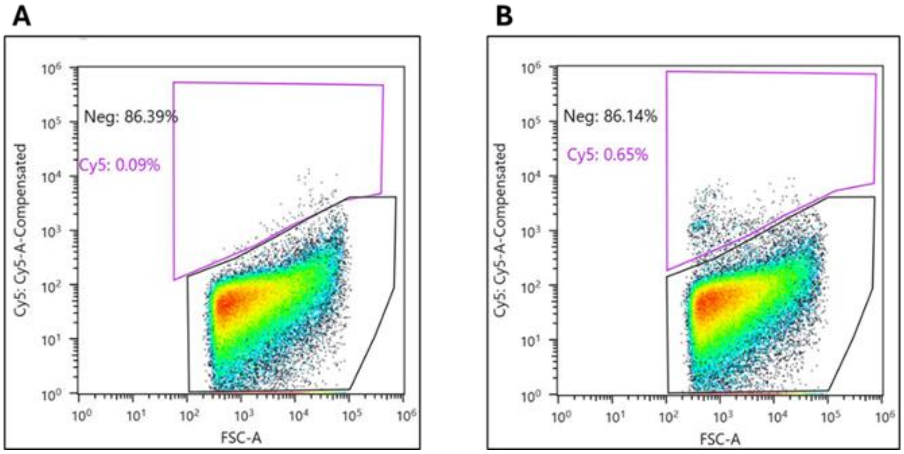


**Figure S7.** Representative FACS plots for 1545B-19F3 No HPG control (A) and 1545B-19F3 Replicate A (B). No-HPG control samples were used to draw the positive (Cy5; purple polygon) and negative (Neg; black polygon) gates for each sample set. The Cy5 positive gate represents BONCAT positive events and is sorted into collection tubes for downstream analysis. The percent values represent the fraction of the total events in each gate.

**Table S1.** Sorting metrics from FACS for each sample. The number of events sorted for each sample and replicate is listed. The percent of total events is the number of positive, sorted events out of the total number of events detected by FACS.

| Sample | Replicate | Sorted BONCAT+ events | Percent of total events |
|---|---|---|---|
| 1546B-1H2 | 1 | 97991 | 0.69% |
| 1546B-1H2 | 2 | 208025 | 1.31% |
| 1546B-1H2 | 3 | 119443 | 0.58% |
| 1545B-19F3 | 1 | 136468 | 0.63% |
| 1545B-19F3 | 2 | 135643 | 0.59% |
| 1545B-19F3 | 3 | 143970 | 0.57% |
| 1547B-1H2 | 1 | 132251 | 0.55% |
| 1547B-1H2 | 2 | 266329 | 0.77% |
| 1547B-1H2 | 3 | 398515 | 0.91% |
| 1549B-3H2 | 1 | 723234 | 6.72% |
| 1549B-3H2 | 2 | 8567 | 0.06% |
| 1549B-3H2 | 3 | 7786 | 0.06% |
| 1548B-3H2 | 1 | 54497 | 0.28% |
| 1548B-3H2 | 2 | 33976 | 0.18% |
| 1548B-3H2 | 3 | 38075 | 0.15% |
| 1547B-8H2 | 1 | 12239 | 0.06% |
| 1547B-8H2 | 2 | 6959 | 0.06% |
| 1547B-8H2 | 3 | 8718 | 0.08% |
