## Supplementary material for "Tracking active heterotrophic microbial communities in the Guaymas Basin deep biosphere with BONCAT-FACS": SI Text

### Supplementary Methods

*Cell Extractions*

Cells were extracted following a modified protocol [1, 2] that we optimized to extract cells from low-biomass subsurface marine sediments. Cell extractions must be adjusted for the particular sample type to maximize cell yield as no single protocol has been demonstrated to work for all sample types (Hatzenpichler 2020). Samples preserved in glycerol-TE were thawed overnight at 4°C and centrifuged at 3,200 × *g* for 30 min to pellet sediment. For each sample, sediment was diluted 1:3 in a filter-sterilized 3.5% NaCl solution (0.2 µm pore size) and briefly homogenized by vortex. From this slurry, 200 µL aliquots were further diluted in 100 µL of pure methanol and 700 µL of filter-sterilized 3.5% NaCl solution. For each experimental replicate, eight extraction replicates were prepared in 2 mL microcentrifuge tubes to ensure enough biomass was extracted for downstream processing. Extraction blanks with phosphate-buffered saline (1X PBS) were processed in parallel to assess contaminant introduction during extraction. Extraction replicates were homogenized by vortex for 30 min at room temperature on setting 7 (approximately 2,000 rpm, VortexGenie2, Scientific Industries, Inc., Bohemia, NY, United States). Cells were separated by density centrifugation by layering 1 mL of 50% Nycodenz solution below the sediment slurry with a syringe and needle and centrifuging at 3,000 × *g* for 10 min. After centrifugation, the upper and inner phases of the supernatant were collected by pipetting and pooling extraction replicates in a 15 mL centrifuge tube. The remaining supernatant was discarded, leaving only the sediment pellet. To detach cells from particles, the pellet was resuspended in 800 µL of 3.5% NaCl solution, 100 µL of pure methanol, and 100 µL of detergent mix (100 mM ethylenediamine tetraacetic acid (EDTA), 100 mM sodium pyrophosphate, 1% Tween80 in 3.5% NaCl solution [1], briefly vortexed to homogenize, and sonicated. The sonication step was done on ice at 125 W and 200 µm amplitude for 5 sec, followed by a 20 sec pause and repeated for 5 cycles to minimize cell rupture. Subsequently, samples were subjected to the same Nycodenz density gradient centrifugation described above. The upper and inner phases of the supernatant from each extraction replicate were combined with the pooled supernatant from the first density gradient centrifugation step. The pooled fractions were centrifuged at 16,000 × *g* for 10 min to pellet extracted cells, and the supernatant was gently decanted, leaving ~1 mL of supernatant remaining to resuspend pelleted cells. This cell suspension was transferred to a 1.5 mL microcentrifuge tube, and cells were centrifuged again at 18,000 × *g* for 10 min and concentrated in 100 µL of PBS:ethanol (1:1 ratio of 1X PBS and pure ethanol). Concentrated cells were stored overnight at -20°C to permeabilize cells for click chemistry.

*Click Chemistry and Fluorescence-Activated Cell Sorting*

The following day, cell extracts were click-stained to fluorescently tag cells that incorporated HPG into newly synthesized proteins [3]. A bulk click solution was prepared, and 200 µL was added to each cell extract. The final click reaction comprised of 500 µM tris-hydroxypropyltriazolylmethylamine (THPTA; Vector Laboratories), 100 µM copper sulfate pentahydrate, 5 mM sodium *L*-ascorbate, 5 mM aminoguanidine HCl, and 5 µM of Cy5 picolyl-azide dye (Vector Laboratories) in 1X PBS. The click solution and cell extract were homogenized and then incubated at room temperature, in the dark, for 30 min. The click solution was washed away with three washes in 1 mL of 1X PBS with centrifugation at 18,000 × *g* for 5 min between each wash. A 50 µL aliquot of the final suspension was kept aside to be used as a "presort" total cell-extractable fraction for PCR. This subsample was stored in a fresh 1.5 mL microcentrifuge tube at -80°C until freeze-thaw DNA extraction.

The remaining cell extract was immediately subjected to fluorescence-activated cell sorting (FACS). Prior to loading, each sample was passed through a 35 µm mesh filter to remove sediment particles and diluted to a final volume of ~1.5 mL with 1X PBS. FACS was performed with a Sony SH800S (Sony Biotechnology, San Jose, CA) equipped with a 70 µm sorting chip using 1X PBS as sheath fluid. The sheath fluid was 0.2 µm filtered and UV sterilized overnight while stirring to remove salt crystals in the PBS and minimize contamination of the FACS fluidics system [4]. A series of gates was manually drawn to exclude large sediment particles. The final sorting gates were drawn based on the Cy5 relative fluorescence and forward scatter (Fig. S7) of the No-HPG controls. A "Cy5-positive" gate was drawn so that only BONCAT-positive cells that incorporated HPG into newly synthesized proteins were sorted. Each sample was sorted until the sample had been exhausted, regardless of the number of events sorted (Table S1). Sorted fractions were centrifuged at 18,000 × *g* for 5 min to pellet cells. After removing the supernatant, cells were resuspended in 50 µL of nuclease-free water and stored at -80°C for freeze-thaw DNA extraction.

*DNA extraction*

DNA from whole sediments was extracted for each replicate sample after HPG incubation using the FastDNA Spin Kit for Soil (MP Biomedicals, Irvine, CA), following the manufacturer's protocol. After elution, DNA was quantified via the Qubit High Sensitivity assay (Invitrogen, Carlsbad, CA) and stored at -80°C until PCR. DNA was extracted from presort and sorted (BONCAT active) fractions by triplicate repeated freeze-thaw cycles to lyse cells [4]. The freeze cycles were done at -80°C for 20 min, and thawing was done at 99°C for 10 min. These DNA extracts were stored at -80°C for further processing.

*16S rRNA Gene Amplicon Sequencing*

Amplification of bacterial and archaeal 16S rRNA genes was performed following the Earth Microbiome protocol [5] using updated primers 515F (forward primer, 5′-GTGYCAGCMGCCGCGGTAA-3’ [6] and 806R (reverse primer, 5′-GGACTACNVGGGTWTCTAAT-3’ [7]. Polymerase chain reaction (PCR) was performed in a final volume of 50 µL consisting of 20 µL of template DNA, 20 µL Invitrogen Platinum Taq II 2x Master Mix, 1 µL of each primer at a concentration of 10 µM, and 8 µL nuclease-free water for freeze/thaw extracted cells, and 25 µL consisting of 5 µL of template DNA, 10 µL Invitrogen Platinum Taq II 2x Master Mix, 0.5 µL of each primer at a concentration of 10 µM, and 9 µL nuclease-free water for DNA extracted with the FastDNA Spin Kit for Soil. The thermocycler conditions were: 94ºC for 3 min, then 28 cycles of 94ºC for 45 sec, 50ºC for 60 sec, and 72ºC for 90ºC, followed by an elongation step of 72ºC for 10 min. A selection of PCR products was checked for the expected length on a 1% agarose gel. All products were then purified using AMPure XP beads following the manufacturer’s protocol with a final elution volume of 35 µL. For Illumina sequencing, Illumina dual index barcodes and sequencing adapters were attached in a second PCR reaction. The final volume for this reaction was 25 µL, consisting of 5 µL of purified amplicon, 10 µL Invitrogen Platinum Taq II 2x Master Mix, 0.5 µL of each primer at a concentration of 10 µM (i5 and i7 primers), and 9 µL nuclease-free water. The thermocycler conditions were: 95ºC for 3 min, then 8 cycles of 95ºC for 30 sec, 55ºC for 30 sec, 72ºC for 30 sec, followed by an elongation step of 72ºC for 5 min. The PCR products were again purified using AMPure XP beads, and then were quantified in triplicate reactions using Quant-iT Picogreen dsDNA assay (Invitrogen) on a Biotek Synergy H1 Hybrid microplate reader. Purified amplicons were then pooled with 7.5 ng of DNA each, or up to 30 μL if a sample had less than 7.5 ng of DNA. The pooled library was quantified with the Qubit high-sensitivity assay. Purification, quality assessment, and sequencing of pooled libraries were performed at the Molecular Research Core Facility at Idaho State University using Illumina's MiSeq chemistry for 2 x 250 bp paired-end reads.

*Bioinformatics Analysis*

Amplicon sequence data were processed with QIIME2 (version 2025.7) [8]. Barcoding and primer sequences were removed using *cutadapt* before truncating the forward (137 bp) and reverse (139 bp) reads using a quality threshold of 30.. DADA2 with default parameters was subsequently used to filter, denoise, and merge the reads. Taxonomic assignments for each ASV (amplicon sequence variant) were determined with the Silva 138.2 database using the *sklearn* method [9]. Contaminant ASVs were identified using the prevalence model (threshold = 0.5) in the R package *decontam* [10] and manual inspection of the dataset to compare samples to the no HPG controls and process blanks. Observed diversity and Shannon's diversity metrics were calculated using the *phyloseq* package in R [11], with replicate samples pooled for analysis. Linear fold enrichment was calculated in Equation 1 as the ratio of the fractional relative abundance of ASVs affiliated with a given class between the presort and sorted fractions:

$$Fold\ Enrichment\ = \left(\frac{max(RAsort, RApresort)\ +0.1}{min(RAsort, RApresort)\ +0.1} - 1\right)\ * sgn(RAsort - RApresort) \qquad (1)$$

Where $RA_{sort}$ and $RA_{presort}$ are the fractional relative abundances of a given taxonomic class in the sorted and presort fractions, respectively. Fold enrichment is calculated in the first part of the function, and a small value (0.1) is added to each relative abundance to avoid dividing by zero. Subtracting that value by one shifts the ratio so that equal relative abundances give zero. The *sgn* function returns the sign of the differences to make fold enrichment favoring the sorted fraction positive, and the fold enrichment favoring the presorted fraction negative.
